# Unmasking *Vibrio paracholerae*: Genomic Reclassification and Epidemiology

**DOI:** 10.1101/2025.09.08.674929

**Authors:** Sergio Mascarenhas Morgado, Erica Lourenço da Fonseca, Ana Carolina Paulo Vicente

**Author notes:** Present address: Av. Brasil, 4365 - Manguinhos, Rio de Janeiro, Brazil.

## Abstract

The genus *Vibrio* encompasses globally relevant pathogens, of which *Vibrio cholerae* is the best known due to its role in cholera. Closely related species within the Cholerae clade—*Vibrio paracholerae, Vibrio metoecus*, and *Vibrio tarriae*—were long misclassified as non-O1/O139 *V. cholerae*. The objective of this study was to analyze all 13,000+ available *V. cholerae* genomes in GenBank to determine the presence of species from the Cholerae clade. Genome-wide analyses using Mash, Whole-genome based Average Nucleotide Identity (gANI), and digital DNA-DNA hybridization (dDDH) reclassified 193 unique genomes as *V. paracholerae*, while *V. metoecus* and *V. tarriae* were not identified. Phylogenomic analyses revealed that *V. paracholerae* forms distinct but relatively homogeneous lineages, spanning clinical, environmental, and animal sources over a period of more than a century. Virulence profiling revealed the absence of CTX and TCP; however, most genomes exhibited other virulence factors, including hemolysins, RTX toxins, colix toxin, and a conserved type VI secretion system. Resistome analysis revealed several antibiotic resistance genes, many of which were incorporated into superintegrons, highlighting the role of *V. paracholerae* as a reservoir of resistance determinants. Importantly, two genomic markers exhibited high discriminatory power for species identification, providing robust tools for epidemiological surveillance. These findings underscore the relevance of *V. paracholerae* for clinical and epidemiological monitoring.

## Introduction

The genus *Vibrio* encompasses globally relevant species that are common etiological agents of diseases in humans and aquatic organisms, with *Vibrio cholerae* being the most well-known as the causative agent of cholera (Baker-Austin et al., 2024). The pathogenicity of certain *V. cholerae* lineages is largely attributable to specific virulence determinants, including the cholera toxin (CTX) and the toxin-coregulated pilus (TCP) (Kumar et al., 2020), which have also been sporadically reported in other *Vibrio* species (Morgado et al., 2024).

In recent years, some species closely related to *V. cholerae* have been identified, most of which were initially misclassified as non-O1/O139 *V. cholerae* (NOVC). These members of the Cholerae clade include *Vibrio paracholerae* (Islam et al., 2021), *Vibrio metoecus* (Kirchberger et al., 2014), and *Vibrio tarriae* (Islam et al., 2022). Although the pathogenic potential of these *V. cholerae*-related species varies, they have also been associated with human infections (Orata et al., 2022). These species appear to coexist with both environmental and clinical *V. cholerae* lineages, some of which are linked to epidemics and pandemics and harbor major cholera virulence determinants such as CTX and VPI-1 (Lemopoulos et al., 2024). This ecological overlap provides an ideal context for the exchange of genetic material, including genes encoding virulence factors (Orata et al., 2015; Islam et al., 2021; Morgado et al., 2024), thereby facilitating the emergence of toxigenic *Vibrio*.

Specifically, *V. paracholerae* has been described as the closest known sister species of *V. cholerae*, based on genetic and genomic analyses (Islam et al., 2021), and has historically been associated with human infections, with reports dating back to 1916 (Dorman et al., 2019). This species has been linked to diarrhea, bacteremia, and sepsis (Morgado et al., 2025). Consequently, its misidentification may introduce bias into epidemiological analyses and hinder appropriate treatment and disease control strategies.

In this study, we conducted a comprehensive genomic and phylogenetic reassessment to clarify the boundaries between *V. cholerae* and other closely related *Vibrio* species, particularly *V. paracholerae*. Using phylogenomics, digital DNA-DNA hybridization (dDDH), and whole-genome Average Nucleotide Identity (gANI), we established reliable cutoffs for species delineation and proposed potential molecular markers.

## Materials and Methods

### Genome Collection and Initial Screening

A total of 13,206 *V. cholerae*, 69 *V. metoecus*, 62 *V. paracholerae*, and 30 *V. tarriae* genomes were retrieved from GenBank (accessed April 2025). To identify potential *V. cholerae* genomes that may have been misclassified, we selected representative reference genomes from closely related species—*V. metoecus* ZF102, *V. paracholerae* NCTC 30, and *V. tarriae* 2521-89. Pairwise genomic distances were estimated using Mash v2.3, which applies a MinHash sketching approach to approximate average nucleotide identity (ANI) (Ondov et al., 2016). Mash distances were calculated between each reference genome and the 13,206 *V. cholerae* assemblies. Genomes presenting higher shared-hash values (i.e., lower Mash distances) with *V. metoecus, V. paracholerae*, or *V. tarriae* references were flagged as candidates for potential misclassification. These genomes were retained for subsequent high-resolution analyses, including gANI and digital DNA–DNA hybridization (dDDH), to confirm species boundaries and refine taxonomic assignments.

### Genomic Similarity and Phylogenetic Analyses

Genomes identified in the Mash-based screening as potential misclassifications were subjected to high-resolution similarity and phylogenetic analyses. Whole-genome average nucleotide identity (gANI) was computed using MiSI (Microbial Species Identifier) (Varghese et al., 2015), which applies genome-wide pairwise BLAST-based comparisons and considers a threshold of ≥96.5% as the cutoff for species delineation. In parallel, digital DNA–DNA hybridization (dDDH) values were estimated using the Genome-to-Genome Distance Calculator (GGDC 3.0) with the recommended *formula 2* (identities/HSP length), which provides robust correlation with traditional wet-lab DNA–DNA hybridization (Meier-Kolthoff et al., 2022).

For phylogenomic reconstruction, we first identified core genes across the dataset using Roary v3.13.0 (Page et al., 2015), with a minimum sequence identity of 95% for orthologous clustering. The resulting core gene alignment was processed with snp-sites v2.5.1 (https://github.com/sanger-pathogens/snp-sites) to extract informative single nucleotide polymorphisms (SNPs). Maximum-likelihood phylogenetic trees were then inferred with IQ-TREE v2.4.0 (Minh et al., 2020), employing the best-fitting nucleotide substitution model selected by ModelFinder and 1,000 ultrafast bootstrap replicates to assess nodal support. Final trees were visualized and annotated using iTOL v6 (Letunic & Bork, 2021).

### Detection of Resistance and Virulence Genes

Antibiotic resistance genes (ARGs) and virulence-associated genes were screened using the abricate tool (https://github.com/tseemann/abricate). Searches were performed against the Comprehensive Antibiotic Resistance Database (CARD) for ARGs and the Virulence Factor Database (VFDB, core dataset) for virulence factors. A minimum threshold of ≥80% sequence identity and ≥80% coverage was applied for gene detection.

To investigate the genomic context of resistance determinants, we further examined the flanking regions of each identified ARG. Specifically, 500 bp of sequences were extracted from both upstream and downstream regions and searched for Vibrio chromosomal repeats (VCRs) using BLASTn. The search considered sequences with >50% identity and >70% alignment coverage.

## Results and Discussion

### Reclassification of *V. cholerae* genomes as *V. paracholerae*

To assess potential misclassified *V. cholerae* genomes, we retrieved all available assemblies from GenBank (n=13,206) and compared them with representative genomes from other members of the Cholerae clade. Pairwise genomic distances were first estimated using Mash, with *V. paracholerae, V. metoecus*, and *V. tarriae* reference genomes as queries against the *V. cholerae* dataset. Genomes exhibiting the highest shared-hashes (≥100/1000) were retained for further analysis using whole-genome average nucleotide identity (gANI). No *V. cholerae* genome exceeded the gANI threshold for species-level assignment to either *V. metoecus* or *V. tarriae*. However, several candidate genomes displayed gANI values consistent with *V. paracholerae*, and were therefore selected for more detailed comparative analyses.

Using *V. paracholerae* NCTC 30 as reference, we identified a preliminary subset of 523/13,206 *V. cholerae* genomes that shared ≥330/1,000 hashes in the Mash analysis, and retained them for downstream comparison. As a baseline, we first calculated gANI values for the 32 genomes previously recognized as *V. paracholerae*, using *V. cholerae* N16961 as reference. These genomes showed gANI values ranging from 95.76% to 96.34% (Table S1), falling below the expected range for species delineation (96.5%). Subsequently, we computed gANI values for the putative set of 523 candidate *V. paracholerae* genomes. Among them, 241 genomes, originally annotated as *V. cholerae*, exceeded the 96.5% gANI threshold, suggesting their reassignment to *V. paracholerae*. Finally, when *V. paracholerae* NCTC 30 was used as the reference, the combined dataset of 273 genomes (32 previously classified as *V. paracholerae* plus the 241 newly identified from the *V. cholerae* dataset) all presented gANI values >96.5%, thereby supporting their classification as *V. paracholerae* (Table S1). Analyses of the 241 candidate “*V. cholerae*” genomes revealed that 48 represented redundant assemblies, which were subsequently removed from downstream analyses, leaving a total of 193 unique genomes (Table S1). These genomes were then subjected to digital DNA–DNA hybridization (dDDH) analysis using *V. cholerae* N16961 as reference. The resulting dDDH values ranged from 64.5% to 68.7% (Table S1), all below the 70% threshold for species delineation, providing further support for their reassignment to *V. paracholerae*.

Finally, the complete set of 225 *V. paracholerae* genomes (32 previously recognized plus 193 reclassified) was subjected to phylogenomic analysis using a core genome MLST (cgMLST) approach. The resulting phylogenetic tree clearly delineated *V. paracholerae* from *V. cholerae* (Figure), confirming that these genomes form a distinct sister clade. These analyses demonstrate that misidentification of *V. paracholerae* as *V. cholerae* occurs in approximately 1.8% of all *V. cholerae* genomes, highlighting a previously underappreciated source of taxonomic and epidemiological bias.

### *V. paracholerae* genomic epidemiology

The cgMLST phylogenomic analysis of the 225 *V. paracholerae* genomes revealed that this species is relatively homogeneous, forming several closely related lineages (Figure). Metadata associated with these genomes (when available) indicated their sources as clinical settings (n = 70), environmental samples (n = 59), and animals (n = 28). Interestingly, some lineages encompassed isolates collected from different sources and across wide temporal and geographic ranges. For example, one lineage includes isolates obtained more than a century apart, from Egypt in 1916 (clinical; GCA_900538065.1) and the USA in 2017 (environmental; GCA_003312095.1). Other noteworthy clusters highlight this diversity: GCA_041005155.1 (USA/2019/animal) and GCA_035782075.1 (Brazil/2001/environment); GCA_045016505.1 (Germany/2021/clinical) and GCA_024105925.1 (Algeria/2018/environment); GCA_019780445.1 (Brazil/2008/clinical) and GCA_003311965.1 (USA/2016/clinical); GCA_006803035.1 (Austria/2011/zooplankton) and GCA_019093165 (China/2018/migratory birds). Of particular interest, two *V. paracholerae* genomes (GCA_030710345.1 and GCA_030718785.1) lacking CTX and TCP were isolated in close association with the pandemic AFR10d and AFR10e lineages of *V. cholerae* during the large cholera outbreak that occurred in the Democratic Republic of the Congo (2009–2012) (Lemopoulos et al., 2024). Taken together, these findings demonstrate that *V. paracholerae* lineages are able to persist across space and time, occurring in the environment (sometimes alongside *V. cholerae*), in clinical contexts, and in animal hosts, highlighting their ecological versatility and epidemiological relevance.

### Identification of a Species-Specific Marker Gene for *V. paracholerae*

For epidemiological tracking purposes, it is crucial to establish reliable markers that can discriminate between *V. cholerae* and *V. paracholerae*. A previous study by Islam et al. (2021) identified two genes: *LysR* family transcriptional regulator (WP_001924807.1) and *HAD-IB* family hydrolase (WP_071179638.1); reported to be present in 22 *V. paracholerae* strains and absent from 22 *V. cholerae* strains. However, when we extended this analysis to our dataset (225 *V. paracholerae* and 12,965 *V. cholerae* genomes), we found that four *V. paracholerae* genomes lacked these genes, while thousands of *V. cholerae* genomes harbored them. These findings demonstrate that the previously genes are not species-specific and cannot reliably distinguish the two taxa.

To overcome this limitation, we performed a pangenome analysis of the 225 *V. paracholerae* genomes and systematically screened candidate markers against the *V. cholerae* dataset. This approach revealed two highly discriminatory genes: (i) *slt* (murein transglycosylase; reference SYZ82122.1), present in all 225 *V. paracholerae* genomes (including one genome with two copies), but detected in only 3/12,965 *V. cholerae* genomes; (ii) *egt*B (ergothioneine biosynthesis; reference SYZ82914.1), present in all *V. paracholerae* genomes, and in only two *V. cholerae* genomes with full coverage (14% coverage in three genomes). Both genes exhibited 100% sensitivity (present in all *V. paracholerae*) and >99.99% specificity (nearly absent in *V. cholerae*), underscoring their potential as robust diagnostic markers for accurate discrimination between these two sister species.

### Virulence and Resistance Gene Profiles

All *V. paracholerae* genomes analyzed lacked the major *V. cholerae* virulence determinants, CTX and TCP (Table S2), a feature also reported for another member of the Cholera clade, *V. metoecus* (Orata et al., 2015). However, fragments of the VPI-2 were detected, with a maximum of 39% coverage in 35 genomes. Within these fragments, the sialic acid metabolism cluster (*nan-nag*) was retained in 35 genomes, including the neuraminidase gene (*nanH*) in 32/35 genomes, whereas the type I restriction–modification system was absent. This pattern of modular fragmentation and retention of VPI-2 loci mirrors observations in *V. metoecus*, in which *nan-nag* cluster was proposed to have originated through independent acquisition of VPI-2 islets (Orata et al., 2015). The *tor* operon, involved in anaerobic respiration of trimethylamine N-oxide and known to enhance cholera toxin production in *V. cholerae* (Lee et al., 2012), was also detected in a truncated form: five genomes contained *tor* genes, with two harboring *tor*AC and three harboring *tor*DR. These results indicate that *V. paracholerae* has undergone either partial acquisition or loss of virulence-associated genomic islands, suggesting divergence in ecological adaptation relative to *V. cholerae*.

Despite lacking the major cholera virulence factors, *V. paracholerae* harbors a broad repertoire of virulence-associated genes. The hemolysin (*hlyA*) and thermolabile hemolysin (*tlh*) were nearly ubiquitous (223/225 genomes), as were RTX toxin components (*rtxB, rtxC*, and *rtxD*, present in 218, 220, and 221 genomes, respectively) (Table S2). The cholix toxin (*chxA*) was identified in 94/225 genomes, consistent with previous observations (Islam et al., 2021). The type VI secretion system (T6SS), implicated in interbacterial competition and environmental fitness (Montero et al., 2023), was present in all genomes, whereas the type III secretion system (T3SS) was restricted to only two genomes (Table S2). The T3SS regions in these two genomes displayed ~97% coverage and identity to *V. cholerae* counterparts, suggesting recent horizontal acquisition likely facilitated by niche overlap between the two species. However, unlike T3SS islands in *V. mimicus* and *V. parahaemolyticus*, these loci lacked the *tdh* and *trh* toxins commonly associated with virulence (Mascarenhas Morgado et al., 2025).

Resistome analysis revealed between 4 and 17 antibiotic resistance genes (ARGs) per genome, with a median of five (Table S3). The most prevalent ARGs were *bla*CARB-9 (n=114), *qnr*VC4 (n=24), *qnr*VC5 (n=13), and *bla*CARB-7 (n=15), along with sporadic genes such as *qnr*VC7, *cat*B9, *aph, flo*R, *dfr*A, *sul, qac*, and *tet*(A/C) (Table S3). Given that *V. cholerae* carries non-mobilizing chromosomal platforms known as superintegrons (SIs), which are hypothesized as reservoirs of ARGs (Rowe-Magnus et al., 2002), we explored this aspect for *V. paracholerae*. By analyzing the genomic context of each ARG, we identified flanking VCRs and/or class 4 integrase (*int*I4) signatures in 102/225 genomes. ARGs confirmed within SIs included *bla*CARB-9 (n=87), *bla*CARB-7 (n=9), *qnr*VC4 (n=21), *qnr*VC5 (n=2), *qnr*VC7 (n=3), *cat*B9 (n=4), and *dfr*A6 (n=1) (Figure). Additionally, 20 genomes contained multiple ARGs within the same SI, with combinations such as *bla*CARB plus *qnr*VC variants or *cat*B9, and in some instances, duplicate copies of the same ARG allele (*bla*CARB or *qnr*VC). As previously demonstrated, ARGs embedded in SIs of *V. paracholerae* can be transcriptionally active and functional (Dorman et al., 2019). These findings support the role of *V. paracholerae* as an ancient and persistent reservoir of ARGs, paralleling the evolutionary dynamics observed in *V. cholerae*.

## Conclusion

A subset of genomes previously classified as *V. cholerae* (~1.8%) were reclassified as *V. paracholerae*, underscoring the misidentification of this sister species in both clinical and environmental surveillance. *V. paracholerae* lineages have demonstrated long-term spatiotemporal persistence across diverse ecological niches, including human clinical cases, animals, and natural environments, reflecting their broad adaptability. Although no toxigenic *V. paracholerae* strains were identified, several genomes were linked to human infections, reinforcing their clinical relevance. Importantly, this study identified robust genetic markers with high discriminatory power for distinguishing *V. paracholerae* from *V. cholerae*, offering a valuable tool for species assignment and epidemiological monitoring.

## CRediT authorship contribution statement

Ana Carolina Vicente: Conceptualization, Methodology, Writing - Original Draft, Writing - Review & Editing, Funding acquisition. Sergio Morgado: Methodology, Formal analysis, Writing - Original Draft, Writing - Review & Editing. Érica Fonseca: Writing - Review & Editing. All authors have read and approved the manuscript.

---

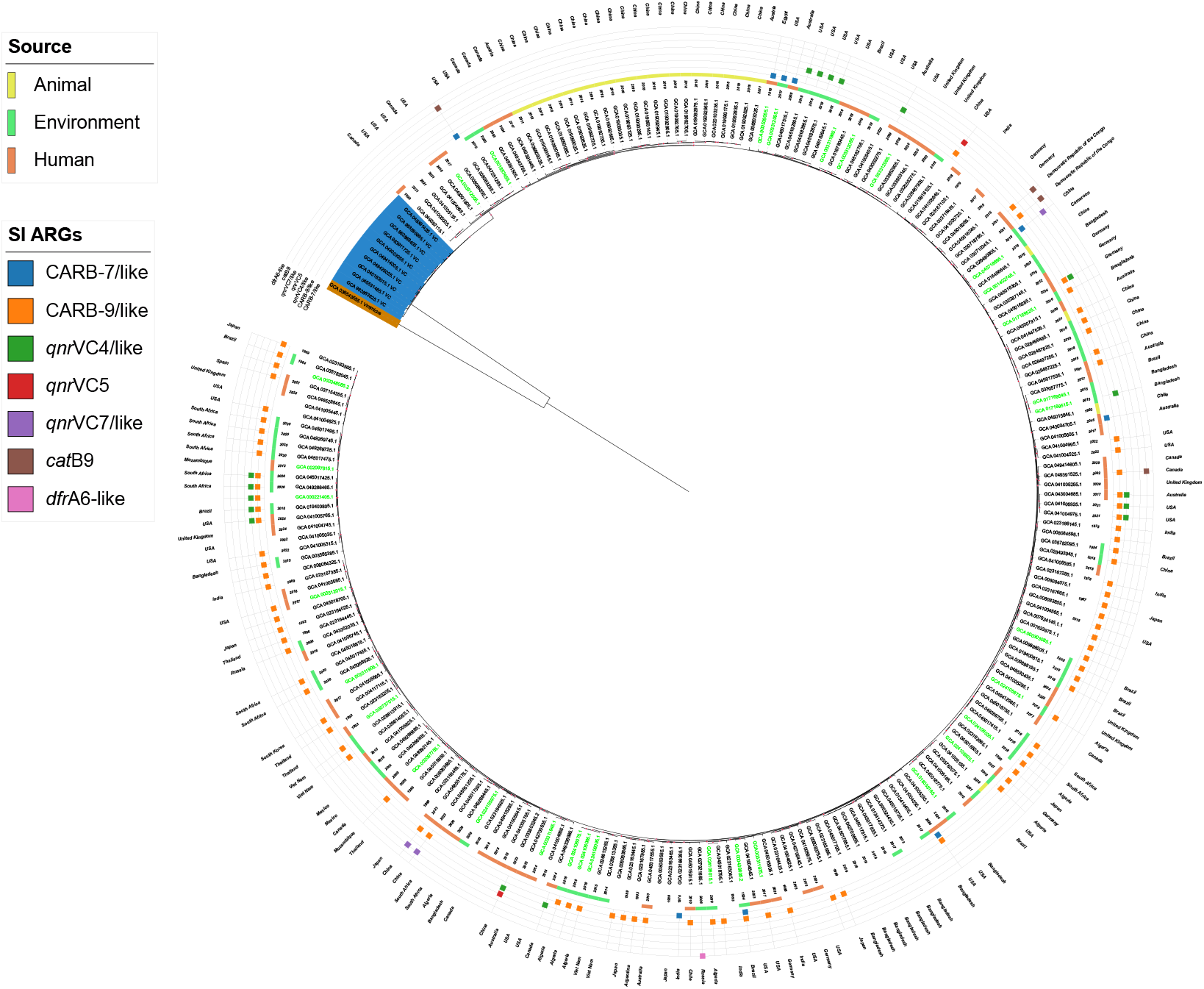

