## Supplementary material for "Unmasking *Vibrio paracholerae*: Genomic Reclassification and Epidemiology": Captions

Figure: Maximum-likelihood cgMLST tree of *Vibrio* genomes. Three Vibrio species are shown: V. mimicus (used as the outgroup, brown background), V. cholerae (blue background), and V. paracholerae. Colored strips adjacent to the labels indicate the source of each genome. Genomes labeled in green represent reference V. paracholerae genomes. Colored squares denote the presence of antibiotic resistance genes within the superintegron of the corresponding genome. Red circles on branches represent >70% bootstrap.
